# Idéfix: identifying accidental sample mix-ups in biobanks using polygenic scores

**DOI:** 10.1101/2021.03.12.435080

**Authors:** Robert Warmerdam, Pauline Lanting, LifeLines Cohort Study, Patrick Deelen, Lude Franke

## Abstract

Identifying sample mix-ups in biobanks is essential to allow the repurposing of genetic data for clinical pharmacogenetics. Pharmacogenetic advice based on the genetic information of another individual is potentially harmful. Existing methods for identifying mix-ups are limited to datasets in which additional omics data (e.g. gene expression) is available. Cohorts lacking such data can only use sex, which can reveal only half of the mix-ups. Here, we describe Idéfix, a method for the identification of accidental sample mix-ups in biobanks using polygenic scores.
In the Lifelines population-based biobank we calculated polygenic scores (PGSs) for 25 traits for 32,786 participants. Idéfix then compares the actual phenotypes to PGSs and uses the relative discordance that is expected for mix-ups, compared to correct samples.

In a simulation, using induced mix-ups, Idéfix reaches an AUC of 0.90 using 25 polygenic scores and sex. This is a substantial improvement over using only sex, which has an AUC of 0.75. Idéfix therefore is not yet able to identify every sample mix-up. However, this will likely improve soon, with highly powered GWAS summary statistics that will likely become available for more commonly measured traits.

Nevertheless, Idéfix can already be used to identify a high-quality set of participants for whom it is very unlikely that they reflect sample mix-ups, and therefore could be offered a pharmacogenetic passport. For instance, when selecting the 10% of participants for whom predicted phenotypes adhere best to the actually measured phenotypes, we estimate that the proportion of sample mix-ups is reduced 250-fold.

**Availability and implementation:** *Idéfix* is freely available at https://github.com/molgenis/systemsgenetics/wiki/Idefix

## 1 Introduction

Biobanks systematically collect (human) biological samples and associated data for research purposes. Enrichment of these biobanks is common through measurements on the collected samples, for example by analyzing serum levels of biological compounds or by genotyping. Repurposing of genetic data for clinical use has become of increasing interest over the past decades (National Academies of Sciences, Engineering, and Medicine *et al.*, 2018). Whereas some initiatives make use of existing diagnostic data (Lee *et al.*, 2020), there are great opportunities for clinical repurposing of existing non-diagnostic data generated by biobanks. However, quality control measures to prevent sample mix-ups (or sample swaps) are commonly less stringent because of the research setting this data was generated in. The existence of sample mix-ups can result in the reporting of results that don’t correspond with the individual and could thereby harm this individual’s health (Gasche *et al.*, 2004)(Ciszkowski *et al.*, 2009)(Heemskerk-Gerritsen *et al.*, 2019). The number of sample swaps that are expected in a research setting varies, with an average having been reported of 3% in genomics datasets (Westra *et al.*, 2011), whereas the frequency of misidentification in laboratory diagnostics is estimated to be between 0.01% and 0.1% (Lippi *et al.*, 2017).

Identifying sample mix-ups in biobank data is a prerequisite to allow it to be used clinically and helps to improve the quality of the data for scientific analysis which will benefit from improved statistical power. It is for example known from simulations that sample mix-ups negatively affect the power of genome-wide association studies (GWASs) and thereby hinder the detection of variants with smaller effect sizes (Zheng and Tian, 2005)(Samuels *et al.*, 2009)(Buyske *et al.*, 2009)(Ho and Lange, 2010).

The most basic test for sample mix-up identification consists of a comparison of the reported sex of individuals and the inferred sex based on the genetic data of these individuals. A drawback of this sex correspondence check is that it is unable to detect sample mix-ups between samples of the same sex. Furthermore, the sex concordance check is only applicable to datasets containing both males and females, and is only able to identify up to 50% of mix-ups given that the dataset contains an equal number of males and females. Additionally, pedigree information can be utilized if family members are included in the biobank and the family relationships are known. A variety of more complex methods have been developed in the past that are able to detect and potentially correct sample mix-ups reliably with the presence of omics datasets (Westra *et al.*, 2011)(Du *et al.*, 2017)(Jiang *et al.*, 2020). As with the sex correspondence check, these methods rely on determining whether or not a phenotype, for example the expression of genes, corresponds to the expected phenotype based on the individual’s genotype. An observed mismatch adds evidence for the sample being a mix-up. Drawbacks of the latter methods are that they either require specific data (such as gene expression or methylation data) that are not commonly available.

In recent years, GWASs have been increasingly powerful, aiding the power and reliability of polygenic scores (PGSs) (Dudbridge, 2013). These PGSs can be used to predict the phenotypes of individuals even if they were not part of the original GWAS. Since PGSs represent an individual’s propensity to a phenotype based on their genetic makeup, they may be useful for identifying sample mix-ups as these PGSs provide an additional method for determining whether or not the sample’s phenotype corresponds to the sample’s genotype. Here, we describe a PGS-based sample mix-up correction method (*Idéfix*) that combines PGSs from multiple traits and determines per sample whether the predicted phenotypes conform to the observed phenotypes, enabling the identification of sample mix-ups. We show our method has predictive power for differentiating between correct samples and sample mix-ups, and that through performing such quantification for a sufficiently number of phenotypes we can reliably identify and remove sample mix-ups.

### 2 Materials and methods

#### 2.1 Datasets

We used adult samples from the Lifelines prospective follow-up biobank (Stolk *et al.*, 2008). These samples were genotyped using the Infinium Global Screening Array® (GSA) MultiEthnic Disease Version 1.0. Within these samples we developed and implemented a method to identify sample mix-ups. To do so, we selected 25 traits for which large-scale GWASs have previously been performed, for which the accompanying summary statistics are available and for which the specific phenotypes were measured in Lifelines. The selected traits and corresponding GWASs are listed in *supplementary table 1.*

#### 2.2 Calculating PGSs

For developing and executing the mix-up identification method PGSs had to be calculated first. A variety of methods and algorithms have been developed over the past years. A recent addition to this range of methods on polygenic prediction is PRS-CS, a tool with superior reported accuracy that functions by inferring posterior effect sizes of single nucleotide polymorphisms (SNPs)(Ge *et al.,* 2019). The method has been shown to equal or outperform competing methods with various GWASs (Chun *et al.*, 2020). Therefore, this tool was chosen to calculate PGSs. GWAS summary statics were processed to comply with the PRS-CS input format. For this we added reference SNP identifiers (RSIDs) to the summary statistics when these were not initially present. This was done by matching the genomic locations of variants to variants from dbSNP (build 137).

The genetic data was preprocessed using PLINK 2.0 (Chang *et al.*, 2015; Purcell and Chang), by excluding ambiguous SNPs and converting the genotype data from VCF to the hybrid PLINK 2.0 bpgen format, maintaining allelic dosage information. Variants were removed if either their minor allele frequency (MAF) was less than 0.01, imputation score was below 0.3, or missing call rates exceeded 0.25. Subsequently, PRS-CS was run to calculate effect sizes given the GWAS summary statistics. We used the European reference linkage disequilibrium (LD) panel provided by the PRS-CS authors. Other parameter settings were left as default. PLINK 2.0 was used to sum variant dosages weighted by the posterior effect sizes as calculated by PRS-CS. The performance of PGSs for continuous traits were assessed by calculating the proportion of the variance in actual phenotypes that is explained by these PGSs (*R*^2^). For ordinal or binary phenotypes, we calculated the area under the ROC curve (AUC). For testing the performance of PGSs all phenotypes were corrected for age, sex and their interaction effects. *R*^2^-values were compared to those reported by literature (*supplementary table 2*).

#### 2.3 Processing of phenotypes

The phenotypes were processed to match the design of the corresponding GWAS, for example exclusion of samples with certain comorbidities or log-transformation. Exclusion criteria and transformations applied are presented in *Supplementary table 3.* Estimated glomerular filtration rate (eGFR) was calculated with the CKD-EPI (Levey *et al.*, 2009). Coronary artery disease was defined as having self-reported balloon angioplasty, bypass surgery or heart attack. Self-reported level of education of Lifelines participants was converted to the number of years of US schooling according to the mapping in *Supplementary table 4* (Okbay *et al.*, 2016). The characteristics of the processed study population is presented in *supplementary table 5*.

#### 2.4 Identifying sample mix-ups

Sample mix-up procedures rely on the (lack of) concordance of an individual’s measured phenotype with their predicted phenotype based on the individual’s genotype. Our method extends this concept by using PGSs to predict an individual’s phenotype. The calculated PGSs are expected to be reasonably accurate for traits like height, where a larger than expected deviance of the predicted phenotype from the measured phenotype adds evidence for a sample being mixed-up. Since the discordance between predicted and measured phenotypes is expected to be relatively high for mix-ups, and relatively low for correct samples, adding additional traits will add predictive power to identify sample mix-ups.

An overview of our method is illustrated in *Figure 1*. The first steps are performed separately for every included trait. As indicated in *Figure 1A,* initially, the relationship between PGSs and actual phenotypes are modeled for the samples that are considered to be correct. For binary traits, like red hair color or the presence of a medical condition, this relationship is modeled using logistic regression. An ordered logistic regression is applied to ordinal traits, which for our set of phenotypes is limited to the spectrum of hair colors ranging from black through brown to blonde.

**Figure 1.**
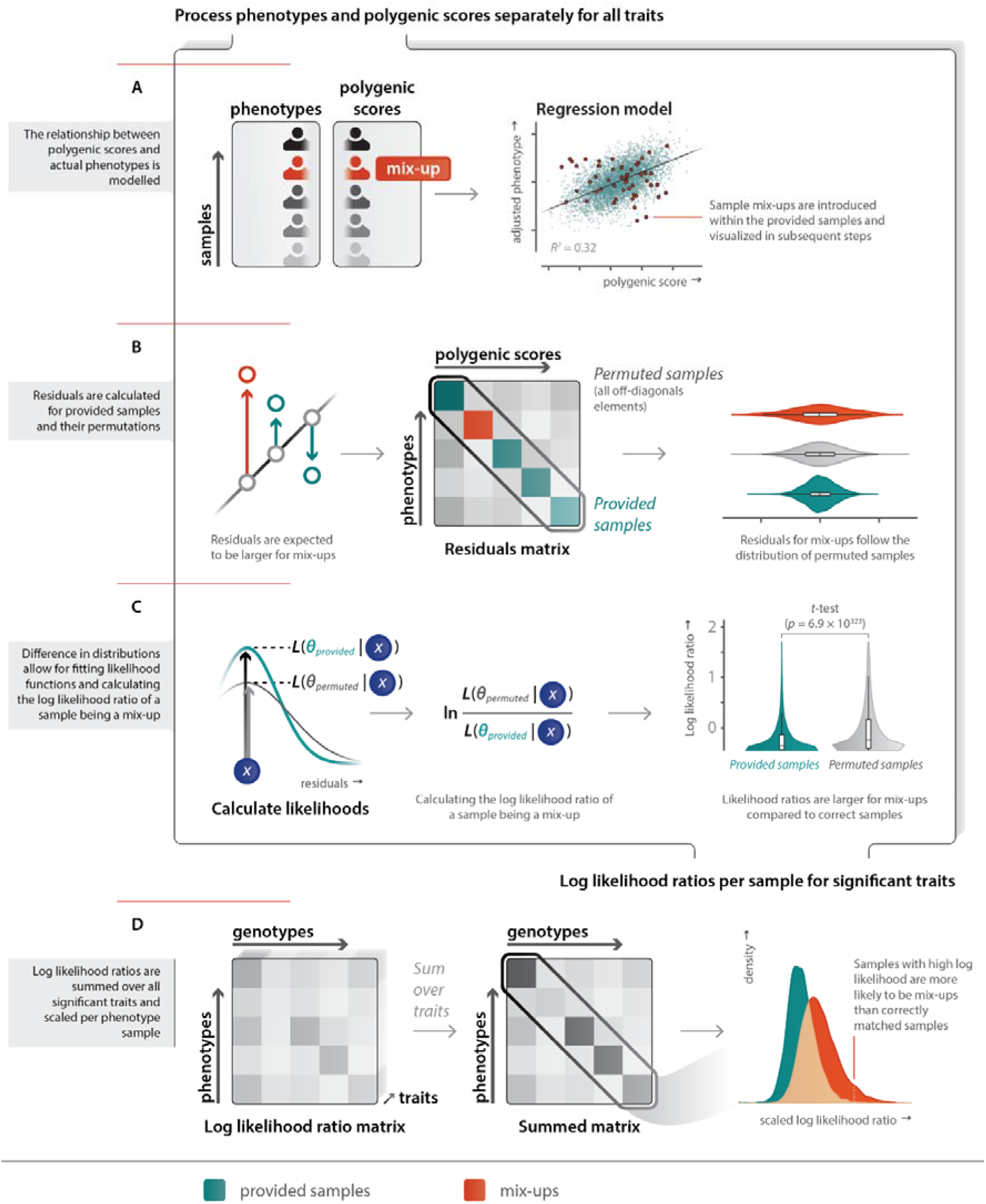
Method overview of that indicates how PGSs and measured phenotypes are used to identify mix-ups. The steps A, B and C are performed separately for all traits. The scatterplot in A, and the distributions visualized in B, C and D are generated using a subset of 5120 samples from the Lifelines dataset of which 1% was introduced as a fake sample mix-up (shown in red). A: The relationship between the input variables is modelled together in a linear model for a continuous trait. This is shown on the right. Introduced fake mix-ups are shown in red. B: Residuals are calculated using the previously fitted model for both the provided sample mappings (main diagonal of the plotted matrix) and the permuted samples (off diagonal in the plotted matrix). The violin plots on the right indicate that permuted samples (grey) and mix-ups (red) are similarly distributed, and differ from the residuals for the provided sample mappings (green). C: (left) For continuous traits gaussian functions are fitted to the permuted (grey) and provided sample mappings (green) to calculate the likelihood of a residual fitting better in the correct or mix-up residual distributions. (middle) Dividing the likelihoods and log-transforming the results in log likelihood ratios of a sample being a mix-up (LLRs). (right) A t-test is employed to test if there is a significant difference between LLRs for permuted and provided sample mappings. D: The matrices on the left and middle indicate summing LLRs over significant traits, and that this aids the predictive power of LLRs. The densities on the right indicate the predictive power of LLRs scaled per row of the LLR matrix.

For quantitative traits, a linear model is used. The summary statistics that are used for calculating PGSs are not able to explain all variation in the phenotype. This is partly due to non-genetic effects. Another cause is that in GWASs, variables that covary with phenotypes, such as age and sex, are often corrected for. The effects of these covariates therefore are not reflected in the summary statistics. To account for the remaining variation that the summary statistics are not able to explain, the predicted phenotypes in our method are modeled using age, sex, and their interaction effect as covariates.

In the following step, for every sample the residuals are calculated, both for the samples that are assumed to be correct – given the provided sample mapping – and their permutations. Here the residuals are essentially the phenotypic variance that cannot be explained by the PGS of a sample. It is therefore expected that the permuted samples have higher residuals compared to correct phenotype-genotype mappings. Permuted samples consist of all other combinations of genotypes and phenotypes, as indicated in *Figure 1B*. In this figure, the different sample mappings are illustrated in a matrix with the measured phenotype part of the samples across the rows, and genotype, or PGS, part of the sample mappings across the columns. The main diagonal herein represents the provided sample mappings, with the off-diagonal representing permuted sample mappings. The figure also shows that the residuals of the provided sample mappings are differently distributed compared to those of both the provided sample mappings and the introduced fake mix-ups.

In case a linear model was employed to model the relationship between the polygenic scores, covariates and the measured phenotypes, residuals are calculated as the deviations of the measured phenotypes from the fitted function. For a logistic regression model, deviance residuals are used to represent residuals which are calculated according to *Equation 1*. Herein *d_i_*, is defined as the deviance residual, *Y_i_* represents the observed outcome and *P_i_* represents the model’s predicted probability for the observed outcome. In case an ordinal logistic regression model is employed, an adaptation of this formula is used to deal with the increase in number of categories.

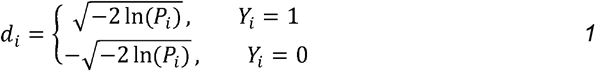

We leverage the different distributions of residuals to calculate log likelihood ratios (LLRs) that indicate whether a sample is likely to be a mix-up or not. These LLRs are calculated on the ratio of the likelihood of a residual fitting in the distribution of provided sample mappings and the likelihood of a residual fitting in the distribution of permuted sample mappings. This step is illustrated in *Figure 1C*. Since residuals are expected to be normally distributed for continuous traits, likelihoods can be calculated by using densities of Gaussian functions fitted to both of the residual distributions. However, this does assume these continuous traits are normally distributed as well, which might not hold for each trait. We therefore ascertained how the performance of this method compared to both an *equal width interval-discretization* and *equal frequency interval-discretization* technique with varying numbers of samples per bin.

##### Equal width interval-discretization

was implemented to separate the parameter space corresponding to the residuals for the provided samples in bins so that each bin corresponds to an interval of equal width. The bins residing at both upper and lower limits of the parameter space are forced to include a minimum number of samples to prevent the model from overfitting, and suffering from outliers having a large effect on separating residuals into bins. This minimum number of samples is also used to prune other bins that do not comply with this lower boundary. Pruned bins are split at their midpoint with the two halves joined to their respective nearest bin. *Equal frequency interval-discretization* was implemented to separate the parameter space corresponding to the residuals for the provided samples in bins so that each bin consisted of an equal number of values.

For both discretization techniques, the breaks obtained within the set of provided samples are subsequently applied to the residuals of permuted samples.

Discretization techniques and the Gaussian likelihood method were assessed by sampling 10,080 samples from the Lifelines dataset and artificially inducing 1% of mix-ups in these subsets. For each subset we applied each discretization technique using 20, 30, 50 and 80 average samples per bin. For the *equal width interval-discretization* the minimum number of samples per bin was set to 10. Corresponding LLRs can be represented as a matrix wherein rows represent phenotype parts of the sample mappings, and columns represent the genotype (PGS) part of the sample mappings. Within this matrix, LLRs were scaled over all rows, or phenotypes. We executed this procedure for 50 individual subsets. Analyses of variance (ANOVAs), followed by a number of Tukey’s honestly significant difference tests indicated that for continuous traits, the gaussian likelihood method outperformed other methods in which a discretization approach was employed *(Supplementary figure 1).*

Phenotypes of binary and ordinal traits are by definition not continuous, and residuals are not normally distributed as is the case for continuous traits. Therefore, using a gaussian function for calculating likelihoods is inappropriate, and the *equal width interval-discretization* technique with an average of 80 samples per bin is used instead. Since we were not able to identify a significant advantage for the binary and ordinal traits for one of the likelihood models, this equal width interval-discretization was chosen over other discretization techniques because it should be less prone to overfitting due to the larger average sample size. Log likelihood ratios are calculated separately for each observed category within an ordinal trait.

We select the traits with significant predictive power for sample mix-ups using a *t*-test to determine whether LLRs of the provided sample mappings differ significantly from those of the permuted sample mappings per trait. In the final step, LLRs are summed over all traits that have a significant difference in the LLRs, resulting in a final matrix of LLRs. Summed LLRs are scaled per phenotype. The threshold for the summed LLRs used to identify sample mix-ups should be determined based on the subsequent use of the resulting dataset. In the following section we highlight a method for determining how an appropriate threshold can be obtained.

##### 2.5 Calculating predictive power of the sample mix-up method

The measure that we considered for assessing the predictive power of the sample mix-up method is the receiver operating characteristics (ROC) curve and the area under the curve. AUC quantifies the ability of a predictor like the LLRs to discriminate between mix-ups and correct samples as the threshold for this separation is varied. The ROC curve can also be used to determine an appropriate threshold. To establish an accurate ROC curve within the Lifelines dataset, we sampled half of the available samples (16,408) as training data and fitted parameters for the method. Thereafter, we took the remaining samples (16,409) and induced 50% mix-ups to validate our model. We used the model fitted on the training data to predict the mix-ups in the validation data. The LLRs were used to establish a ROC curve and accompanying area under the curve (AUC). From the LLRs and ROC curve we have extracted a number of thresholds. This was compared to the performance of a regular sex correspondence check in which the inferred sex from genotypes was compared to the reported sex to find mismatches. We also looked into the performance of a combined predictor that included both the PGS sample mix-up identification method and the sex correspondence check.

Our method is implemented in the R programming language (version 3.6.1). Ordinal logistic regression models were fitted using the MASS package (version 7.3-51.6) (Venables and Ripley, 2002). ROCs and AUCs were calculated using the pROC package (version 1.16.2) (Robin *et al.*, 2011).

### 3 Results

#### 3.1 Polygenic score-based sample mix-up identification

We have developed a sample mix-up identification method (*Idéfix*) that relies on the comparison of actual phenotypes to PGSs. Our method works by:

- modelling the relationships between phenotypes and polygenic scores,
- calculating the residuals of the provided samples and their permutations,
- employing a likelihood model fitted on the residuals for provided and permuted samples,
- combining likelihood ratios over multiple traits.

We implemented our method in the R programming language (version 3.6.1). The installation and usage instructions for *Idéfix* is described on the wiki. (https://github.com/molgenis/systemsgenetics/wiki/Idefix)

#### 3.2 Polygenic scores

We calculated PGSs for the 25 selected traits in the Lifelines dataset. In order to assess the predictive power of the PGSs for continuous traits *R*^2^-values were calculated. For ordinal or binary phenotypes, we calculated spearman correlation and AUC’s respectively. These values indicate that for most traits, the performance is consistent with prior reported performance in literature. The explained variance ranges from 3.0% for the concentration of basophilic granulocytes to 33.9% for height. However, we observed that PGSs for body mass index (BMI) and educational attainment do not explain as much variance compared to previously reported values (11.9% and 5.3% respectively). The explained variances for continuous traits are presented in *Supplementary figure 2.* This figure also illustrates that employing a model that corrects for both sex and age is, as expected, increase the phenotypic variance that PGSs explain.

#### 3.2 Predictive power of our method

We assessed the ability of *Idéfix* to discriminate between sample mix-ups and correct samples by executing the method we developed in the Lifelines dataset using the PGSs we calculated and the processed phenotypes. We applied the method on half of the samples in which we introduced 50% sample mix-ups, using the regression and likelihood models fitted to the other half of the samples in which we did not introduce artificial mix-ups. The measurements for the performance are therefore also suitable for situations in which a regular, moderate, percentage of mix-ups are expected. This approach allowed us to get an accurate measurement of the performance of our method for individual traits, as well as for the method overall. First of all, we see that AUCs for individual traits ranges from 0.50 to 0.66. In *supplementary figure 3* is presented that for continuous traits, these AUCs are positively correlated with the variance explained by PGSs. The number of traits available for the samples in our dataset is presented in *supplementary table 6.*

In *figure 2* is shown that the AUC for the polygenic score-based sample mix-up identification for the 25 traits combined is 0.80. Using the AUC as a measure for performance, we also notice that our method performs better than the sex correspondence check, although this is able to identify 50% of samples without false positives if an equal number of males and females are present in the dataset. When combining the sex-check with the PGS-based sample mix-up identification the AUC increases to 0.90. It is conceivable that different thresholds are advantageous dependent on the intended application of the genetic data. For instance, GWASs profit from increased sample sizes (Canela-Xandri *et al.*, 2018), making lenient thresholds more suitable for this application compared to more stringent thresholds. Alternatively, when repurposing genetic data for clinical use more stringent thresholds are desired. In *figure 2* we estimate that the proportion of sample mix-ups is reduced 250-fold when selecting the 10% of participants for whom predicted phenotypes adhere best to the measured phenotypes

**Figure 2.**
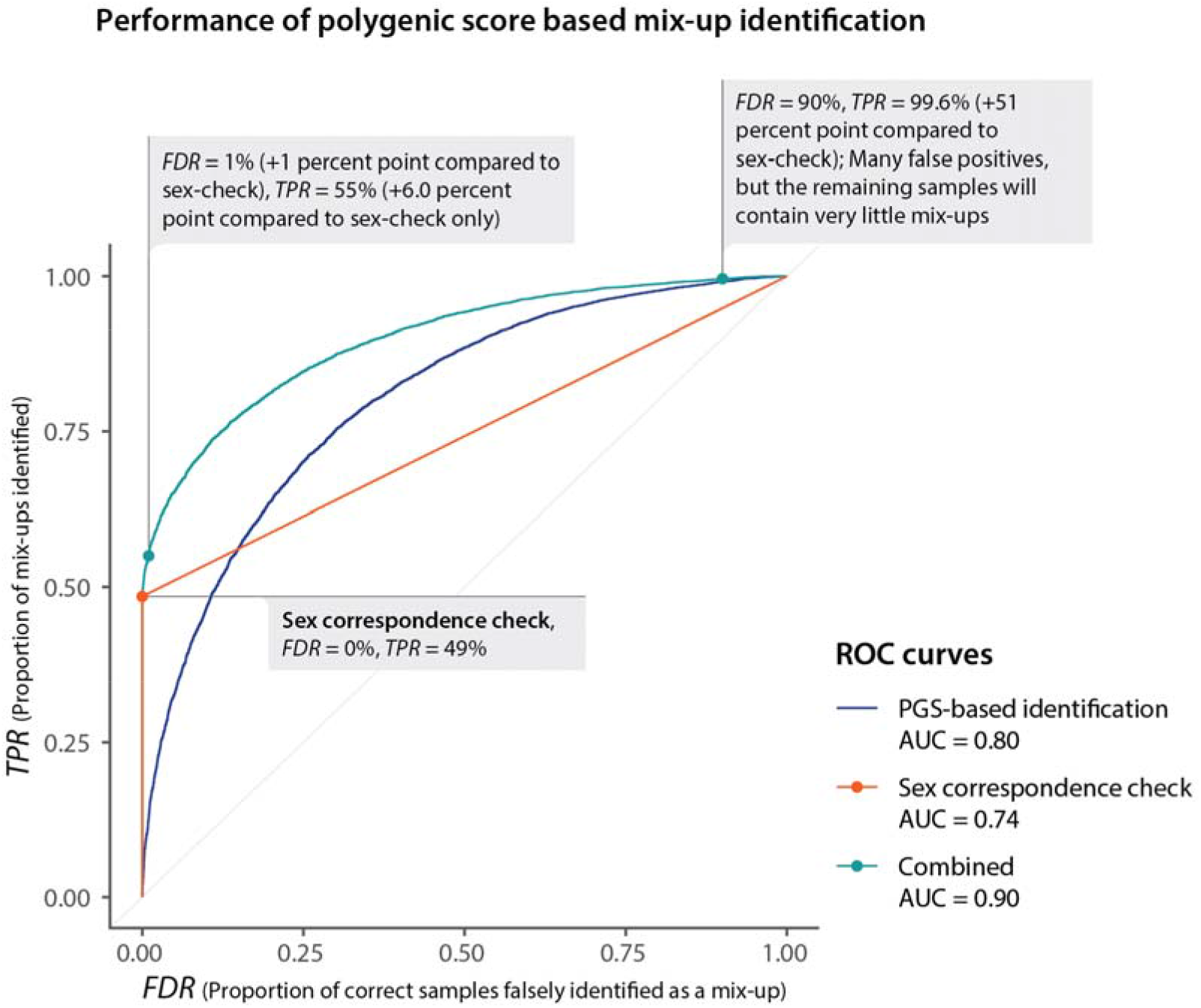
Performance of the polygenic score-based mix-up predictor (blue), the sex concordance check (sex correspondence check, orange), and a combined predictor (green) as illustrated by receiver operating characteristics (ROC). The x-axis indicates the proportion of correct samples that are falsely identified as a mix-up, named the false discovery rate (FDR). The y-axis represents the proportion of mix-ups that are identified as mix-ups, named the true positive rate (TPR). Coordinates of one of the curves correspond to the FDR and TPR for a particular threshold of the predictor. Due to male-female imbalance in the dataset, the proportion of mix-ups identified as shown for the sex correspondence check deviates from the expected value of 0.5. Because of this deviation, the AUC is 0.74 as opposed to the expected AUC of 0.75 given an equal number of males and females.

#### 3.3 application in Lifelines

Applying Idéfix to our entire cohort enabled us to identify a number of potential sample mix-ups. To our surprise, four non-European samples have considerably higher predictions to be mix-ups relative to other samples, despite the fact that 98.2% of the analyzed samples are of Dutch ancestry. To ascertain that these findings are not ethnicity-driven, samples of European and non-European ethnicity were compared. This indicated that for non-European samples, there is on average not more evidence for classifying these as sample mix-ups (Welch’s *t*-test, *p* = 0.92), strengthening the support for the identified samples being mix-ups (supplementary figure 4). By plotting the discordance between actual and predicted phenotypes we can visually inspect how *Idéfix* identifies potential mix-ups (supplementary figure 5), we see that each of our top four predicted mix-up samples show systematic deviation for multiple traits, which indicates that not a single trait is driving the predictions. We also observe large deviations for height (*p*-values between 3.07 × 10^−3^ and 1.13 × 10^−8^). Given these observations, we conclude these are likely mix-ups.

### 4 Discussion

Sample mix-ups present a challenge for the repurposing of genetic data available in biobanks, as well as hindering the capability of association analyses to detect small genetic effects. Existing methods to resolve mix-ups are either not able to identify sample mix-ups within the same sex, or require additional data like gene expression or methylation measurements. For most biobanks such genomics data is usually not available. This was the motivating reason to develop *Idéfix,* a method that is able to identify sample mix-ups in biobanks while solely relying upon genetic data and a limited number of measured phenotypes, such as height, BMI, cholesterol and hair color.

We have shown that our method is capable of identifying sample mix-ups with high accuracy. This is extremely valuable for biobanks that wish to return information to their biobank participants, such as a pharmacogenetic passport or polygenic risk scores for certain diseases that can be (partly) prevented or intervened on by lifestyle modification. In order to do so, it is crucial to know that information that is being returned to individuals is accurate, and our method *Idéfix,* can help to ensure this to be the case.

Previously, a problem regarding sample identification has been described as well (Cai *et al.*, 2017; Lippert *et al.*, 2017), although the performance of one of these methods was not clear (Erlich, 2017). Here, we have leveraged a set of well-powered GWASs, in conjunction with a recent method for calculating PGSs, and a likelihood ratio framework to accurately weigh these PGSs to determine to what extent sample mix-ups can be identified. Most importantly, we developed this method because we aimed to identify a set of samples that very likely do not contain mix-ups, and whom for instance can safely be returned pharmacogenetic passports. Having such a sub-cohort of biobank participants available can, for instance, be very valuable for investigating how biobank participants experience pharmacogenetics.

We do observe several avenues to improve our method. We have shown that our method performs best with PGSs from well powered GWASs *(supplementary figure 3*). However, PGSs do not capture the full heritability of traits. Furthermore, GWASs that identify variants associated with a particular trait which are the basis of PGSs are focused on common variants, while rare variants tend to have larger effect sizes (Marouli *et al.*, 2017). While an increase in sample sizes will probably result in rarer alleles being discovered, the accuracy of PGSs can also be improved by integrating large-effect expression variants found in other ways as well (Smail *et al.*, 2020). Larger sample sizes also increase the power of GWASs to identify variants regardless of their effect size (Canela-Xandri *et al.*, 2018), increasing the subsequent predictive power of PGSs yet more. Another limitation of PGSs is that these are much less predictive for non-European cohorts, supporting the need for large-scale GWASs in diverse human populations (Duncan *et al.*, 2019). A consequence of the less performant PGSs in non-European samples is that there might be less power to detect structural discordances with phenotypes, causing results to be differently distributed for these samples compared to European samples (supplementary figure 4). Although our sample-size did not permit us to find a significant difference, we suggest handling different ethnicities with care. We do expect that this limitation will be resolved when GWAS are performed on mixed ancestry in cosmopolitan cohorts. Until then one possible solution is to run *Idéfix* separately for each population separately.

In the past years numerous biobanks have been established, each of these providing valuable resources on genotype data and a large number of phenotypes. This is a good starting point for our method. However, since the discovery and validation samples should be independent when calculating PGSs (Wray *et al.*, 2013), this poses a problem when biobanks are included in GWASs that are used for calculating PGSs since these will then be biased for the included biobanks. However, we expect that the effect on the performance of *Idéfix* is limited. Since current GWASs are performed on many different cohorts, the effect of each individual or cohort on the PGSs is limited.

Biobanks and studies often differ in the phenotypes that have been measured and that are available. With different phenotypes being available for other biobanks, there is the opportunity for additional heritable phenotypes to be included. However, this variability also limits the applicability to cohorts that have fewer phenotypes available since a limited number of phenotypes will limit the performance of our method, although it will still add to the performance of a sex-based concordance check. Additionally, the first step of the method we developed is dependent in modelling the relationship between the measured phenotypes of individuals and the PGSs. Identifying such a relationship can be hindered by a lack of samples or be biased when there are too many sample mix-ups present in the dataset.

Our method can be beneficial in a variety of cases. For instance, this method is able to identify mix-ups irrespective of the proportion of males and females included in the study, whereas a common concordance check on sex alone is only able to perform optimally with an equal number of individuals for both sexes. Our method relies on the availability of multiple traits in the biobank for which PGSs can be calculated. Furthermore, it is possible for our method to be expanded with additional traits when these become available in the future and it is expected that performance will increase dependent on the heritability of the trait. In the Lifelines cohort, 285 sample mix-ups have already been identified, 42% of which have been identified by utilizing pedigree information and ascertaining whether the genotype data is concordant with these reported familial relationships. These samples could not have been identified using sex-check alone, exemplifying the necessity of additional mix-up identification in biobanks wherein familial relationships cannot be used. Despite the thorough quality control, *Idéfix* has enabled us to identify 4 additional sample mix-ups with high confidence.

Currently, *Idéfix* is not yet able to identify every single sample mix-up: it can happen that for a certain sample the calculated PGS for each of the 25 phenotypes are all quite average and that the observed, but incorrect phenotypic measurements behave similarly. In that case this sample is not yet flagged as a sample mix-up. However, by using larger GWAS studies in the near future that lead to more accurate PGSs and by including more traits we expect our method to help resolve most sample mix-ups in biobanks within the next few years.

## Lifelines Cohort Study

The Lifelines Biobank initiative has been made possible by funding from the Dutch Ministry of Health, Welfare and Sport, the Dutch Ministry of Economic Affairs, the University Medical Center Groningen (UMCG the Netherlands), University of Groningen and the Northern Provinces of the Netherlands. The generation and management of GWAS genotype data for the Lifelines Cohort Study is supported by the UMCG Genetics Lifelines Initiative (UGLI). The authors wish to acknowledge the services of the Lifelines Cohort Study, the contributing research centers delivering data to Lifelines, and all the study participants.

Raul Aguirre-Gamboa (1), Patrick Deelen (1), Lude Franke (1), Jan A Kuivenhoven (2), Esteban A Lopera Maya (1), Ilja M Nolte (3), Serena Sanna (1), Harold Snieder (3), Morris A Swertz (1), Judith M Vonk (3), Cisca Wijmenga (1)

1) Department of Genetics, University of Groningen, University Medical Center Groningen, The Netherlands

2) Department of Pediatrics, University of Groningen, University Medical Center Groningen, The Netherlands

3) Department of Epidemiology, University of Groningen, University Medical Center Groningen, The Netherlands

We thank the UMCG Genomics Coordination Center, the UG Center for Information Technology and their sponsors BBMRI-NL & TarGet for storage and compute infrastructure.

We thank Esteban Lopera-Maya for executing and elucidating the original quality control procedures.

We thank Shai Carmi for his feedback on our preprint.

We thank Albert Uderzo and René Goscinny for inspiring us to call our method Idéfix.

---

*Supplementary figure 1:* A comparison between 9 likelihood models for the 25 traits. For every trait the nine likelihood models were tested 50 times on a randomly selected subset of samples and fake mix-ups. The area under the ROC curve for all combinations is represented by the y-axes. The iterations are combined in a violin plot, with dots and error bars illustrating the mean and standard deviations respectively. The p-values for analyses of variance (ANOVA) are included for every trait. The outcome for Tukey’s HSD pairwise tests with the Gaussian model are shown above the violin plots for the other eight models. (****: *p* ≤ 1 × 10^−4^, ***: *p* ≤ 1 × 10^−3^, **: *p* ≤ 1 × 10^−2^, *: *p* ≤ 0.05, else: *p* > 0.05).

*Supplementary figure 2:* Here we show the importance of first adjusting the data for age, sex and their interaction effects. The variance of phenotypes explained (*R*^2^) by polygenic scores (PGSs) is shown as reported in literature, compared to what we achieved in the Lifelines dataset. A base value (unadjusted) is shown for the Lifelines dataset as well a value calculated using phenotypes that were adjusted for age, sex and their interaction effects. When using such adjustments, *R*^2^-values become comparable to those reported by literature, which are also calculated using similar corrections. For blood traits a male stratified approach was used instead of a correction model. The estimated *R*^2^-value for estimated glomerular filtration rate (eGFR) as reported by literature is calculated using only index SNPs.

*Supplementary figure 3:* The power of polygenic scores (PGSs) for identifying sample mix-ups for individual continuous traits, as a function of the variance of the trait explained by the polygenic scores. The predictor used for the area under the ROCs are log likelihood ratios for a sample being mixed up, scaled within all combinations of individual phenotype samples.

*Supplementary figure 4:* A comparison of predictions between European and non-European samples (based on principal components of genotype data). There is no significant difference between the two distributions (Welch’s *t*-test, *p* = 0.92).

*Supplementary figure 5:* A visualization of the four samples with the highest predictions for being mixed-up compared to four samples with the lowest predictions. The four samples with the highest predictions show larger residuals compared to the four samples with the lowest predictions. The x-axis represents the absolute scaled residuals. Residuals were scaled per phenotype in case the trait is on an ordinal or binary scale. Traits with absolute scaled residuals larger than 2.58 (*p* = 0.01) are labelled with a *p*-value.

*Supplementary table 1:* An overview of the included traits, the corresponding GWASs and the original paper these were originally published in.

*Supplementary table 2:* Collection of *R*^2^-values and AUCs as reported by literature.

*Supplementary table 3:* An overview of the phenotype processing that was applied for the included traits. Age and sex were used in modelling the relationships between polygenic scores and measured phenotypes. Samples for which any of the other conditions applied which are listed in the correction column were excluded from the trait for which the condition was set.

*Supplementary table 4:* A table indicating how answers to the question “What is the highest level of education you have finished?” were mapped to US years of schooling.

*Supplementary table 5:* Study population characteristics. For every trait and category within that trait (if applicable) the characteristics are given for those samples that fall within that group. The columns indicate if the trait, the category (if the trait is categorical), the mean of the ages for samples the row pertains to, the corresponding standard deviation for these ages, the number of males, the number of females, the mean phenotypes for the trait, the standard deviation in the phenotypes for the trait, the number of samples. The last column indicates what percentage of the total number of samples, for which the categorical trait is available, belong to the specified category.

*Supplementary table 6:* The distribution of the number of traits available for ROC estimation.

## References

Broman, K.W. et al. (2015) Identification and Correction of Sample Mix-Ups in Expression Genetic Data: A Case Study. G3amp58 GenesGenomesGenetics, 5, 2177–2186.

Buyske, S. et al. (2009) When a Case Is Not a Case: Effects of Phenotype Misclassification on Power and Sample Size Requirements for the Transmission Disequilibrium Test with Affected Child Trios. Hum. Hered., 67, 287–292.

Cai, B. et al. (2017) Matching phenotypes to whole genomes: Lessons learned from four iterations of the personal genome project community challenges. Hum. Mutat., 38, 1266–1276.

Canela-Xandri, O. et al. (2018) An atlas of genetic associations in UK Biobank. Nat. Genet., 50, 1593–1599.

Chang, C.C. et al. (2015) Second-generation PLINK: rising to the challenge of larger and richer datasets. GigaScience, 4.

Chun, S. et al. (2020) Non-parametric polygenic risk prediction using partitioned GWAS summary statistics. bioRxiv, 370064.

Ciszkowski, C. et al. (2009) Codeine, Ultrarapid-Metabolism Genotype, and Postoperative Death. N. Engl. J. Med., 361, 827–828.

Demenais, F. et al. (2018) Multiancestry association study identifies new asthma risk loci that colocalize with immune-cell enhancer marks. Nat. Genet., 50, 42–53.

Du, Y. et al. (2017) A SNP panel and online tool for checking genotype concordance through comparing QR codes. PLOS ONE, 12, e0182438.

Dudbridge, F. (2013) Power and Predictive Accuracy of Polygenic Risk Scores. PLOS Genet., 9, e1003348.

Duncan, L. et al. (2019) Analysis of polygenic risk score usage and performance in diverse human populations. Nat. Commun., 10, 3328.

Erlich, Y. (2017) Major flaws in “Identification of individuals by trait prediction using whole-genome sequencing data”. bioRxiv, 185330.

Evangelou, E. et al. (2018) Genetic analysis of over 1 million people identifies 535 new loci associated with blood pressure traits. Nat. Genet., 50, 1412–1425.

Gasche, Y. et al. (2004) Codeine Intoxication Associated with Ultrarapid CYP2D6 Metabolism. N. Engl. J. Med., 351, 2827–2831.

Ge, T. et al. (2019) Polygenic prediction via Bayesian regression and continuous shrinkage priors. Nat. Commun., 10, 1–10.

Heemskerk-Gerritsen, B.A.M. et al. (2019) Survival after bilateral risk-reducing mastectomy in healthy BRCA1 and BRCA2 mutation carriers. Breast Cancer Res. Treat., 177, 723–733.

Ho, L.A. and Lange, E.M. (2010) Using public control genotype data to increase power and decrease cost of case–control genetic association studies. Hum. Genet., 128, 597–608.

Hoffmann, T.J. et al. (2018) A large electronic-health-record-based genome-wide study of serum lipids. Nat. Genet., 50, 401–413.

Jiang, Y. et al. (2020) DRAMS: A tool to detect and re-align mixed-up samples for integrative studies of multi-omics data. PLOS Comput. Biol., 16, e1007522.

Lee, J.J. et al. (2018) Gene discovery and polygenic prediction from a genome-wide association study of educational attainment in 1.1 million individuals. Nat. Genet., 50, 1112–1121.

Lee, M. et al. (2020) Repurposing of Diagnostic Whole Exome Sequencing Data of 1,583 Individuals for Clinical Pharmacogenetics. Clin. Pharmacol. Ther., 107, 617–627.

Levey, A.S. et al. (2009) A new equation to estimate glomerular filtration rate. Ann. Intern. Med., 150, 604–612.

Lippert, C. et al. (2017) Identification of individuals by trait prediction using whole-genome sequencing data. Proc. Natl. Acad. Sci.

Lippi, G. et al. (2017) Managing the patient identification crisis in healthcare and laboratory medicine. Clin. Biochem., 50, 562–567.

Mahajan, A. et al. (2018) Fine-mapping type 2 diabetes loci to single-variant resolution using high-density imputation and islet-specific epigenome maps. Nat. Genet., 50, 1505–1513.

Marouli, E. et al. (2017) Rare and low-frequency coding variants alter human adult height. Nature, 542, 186–190.

National Academies of Sciences, Engineering, and Medicine et al. (2018) Returning Individual Research Results to Participants: Guidance for a New Research Paradigm Downey, A.S. et al. (eds) National Academies Press (US), Washington (DC).

Okbay, A. et al. (2016) Genome-wide association study identifies 74 loci associated with educational attainment. Nature, 533, 539–542.

Purcell, S. and Chang, C. PLINK 2.0.

Robin, X. et al. (2011) pROC: an open-source package for R and S+ to analyze and compare ROC curves. BMC Bioinformatics, 12, 77.

Samuels, D.C. et al. (2009) Detecting new neurodegenerative disease genes: does phenotype accuracy limit the horizon? Trends Genet., 25, 486–488.

Smail, C. et al. (2020) Integration of rare large-effect expression variants improves polygenic risk prediction. medRxiv, 2020.12.02.20242990.

Stolk, R.P. et al. (2008) Universal risk factors for multifactorial diseases: LifeLines: a three-generation population-based study. Eur. J. Epidemiol., 23, 67–74.

van der Harst Pim and Verweij Niek (2018) Identification of 64 Novel Genetic Loci Provides an Expanded View on the Genetic Architecture of Coronary Artery Disease. Circ. Res., 122, 433–443.

Venables, W.N. and Ripley, B.D. (2002) Modern Applied Statistics with S Fourth. Springer, New York.

Vuckovic, D. et al. (2020) The Polygenic and Monogenic Basis of Blood Traits and Diseases. Cell, 182, 1214–1231.e11.

Westra, H.-J. et al. (2011) MixupMapper: correcting sample mix-ups in genome-wide datasets increases power to detect small genetic effects. Bioinformatics, 27, 2104–2111.

Wheeler, E.et al. (2017) Impact of common genetic determinants of Hemoglobin A1c on type 2 diabetes risk and diagnosis in ancestrally diverse populations: A transethnic genome-wide meta-analysis. PLoS Med., 14, e1002383.

Wray, N.R. et al. (2013) Pitfalls of predicting complex traits from SNPs. Nat. Rev. Genet., 14, 507–515.

Wuttke, M. et al. (2019) A catalog of genetic loci associated with kidney function from analyses of a million individuals. Nat. Genet., 51, 957–972.

Yengo, L. et al. (2018) Meta-analysis of genome-wide association studies for height and body mass index in ~700000 individuals of European ancestry. Hum. Mol. Genet., 27, 3641–3649.

Zheng, G. and Tian, X. (2005) The impact of diagnostic error on testing genetic association in case-control studies. Stat. Med., 24, 869–882.

