## Supplementary figures and images for "Idéfix: identifying accidental sample mix-ups in biobanks using polygenic scores"

### Supplementary figure 1

#### Comparison of likelihood models

A: 1–15 / 25 traits

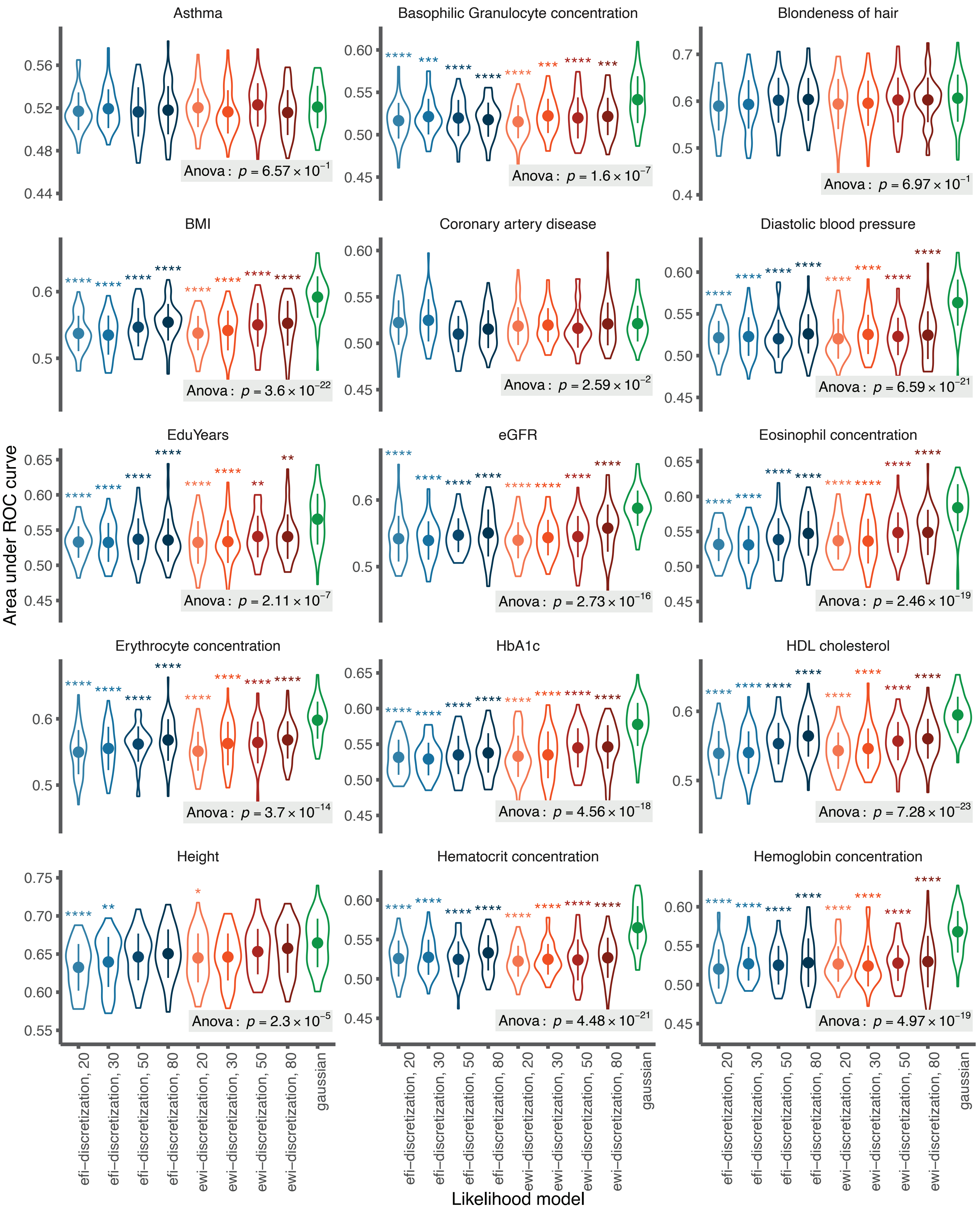

##### Comparison of likelihood models

B: 16–25 / 25 traits

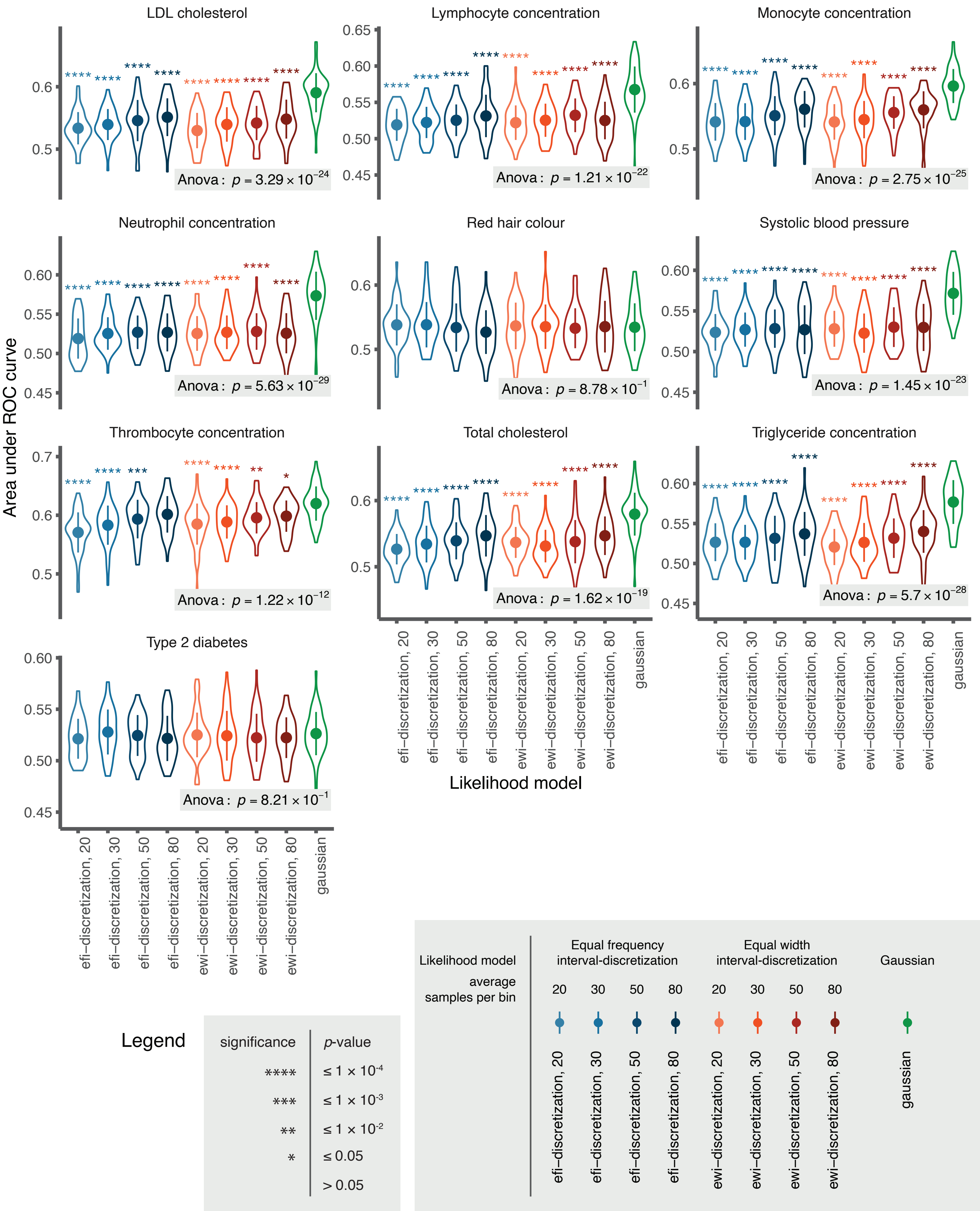

### Supplementary figure 2

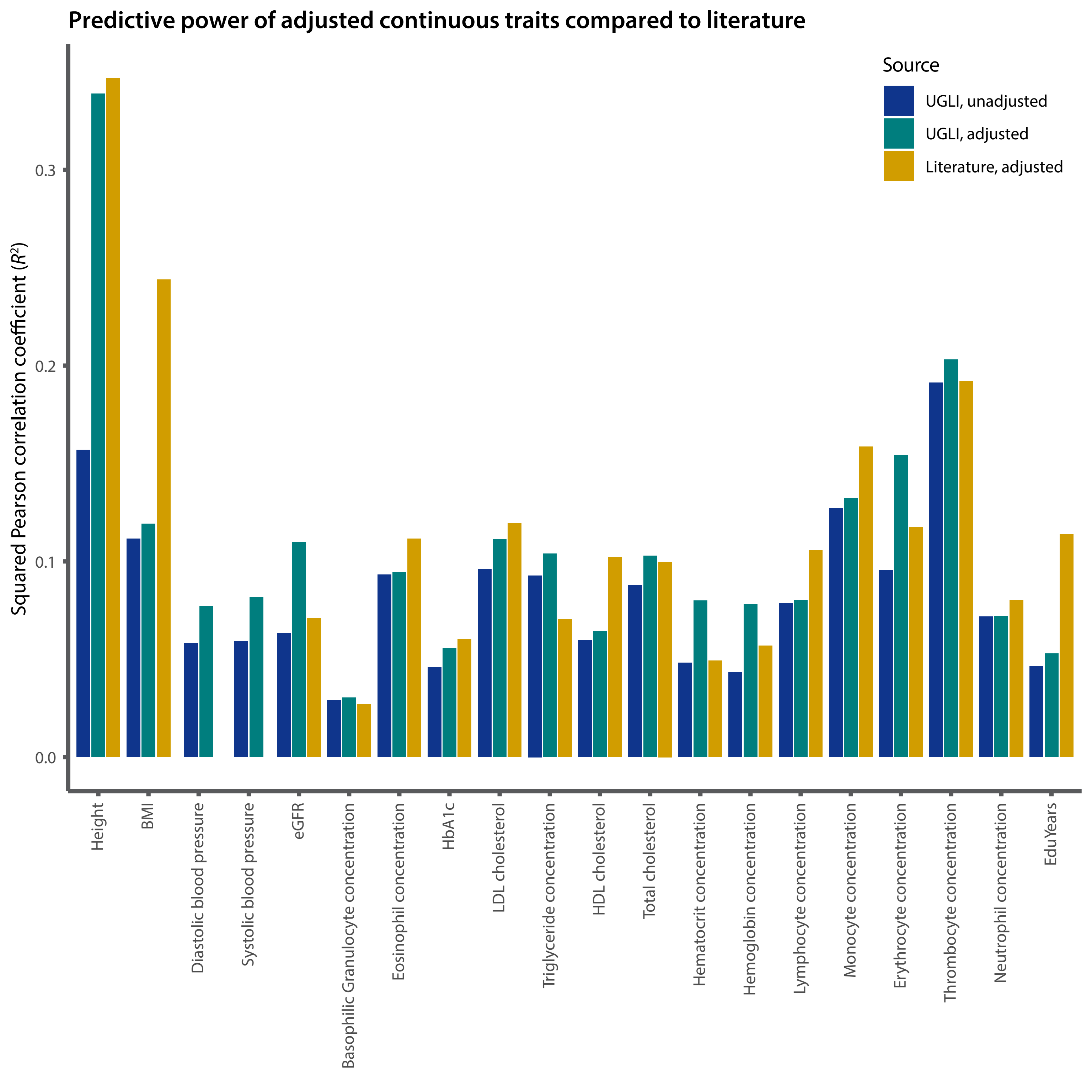

### Supplementary figure 3

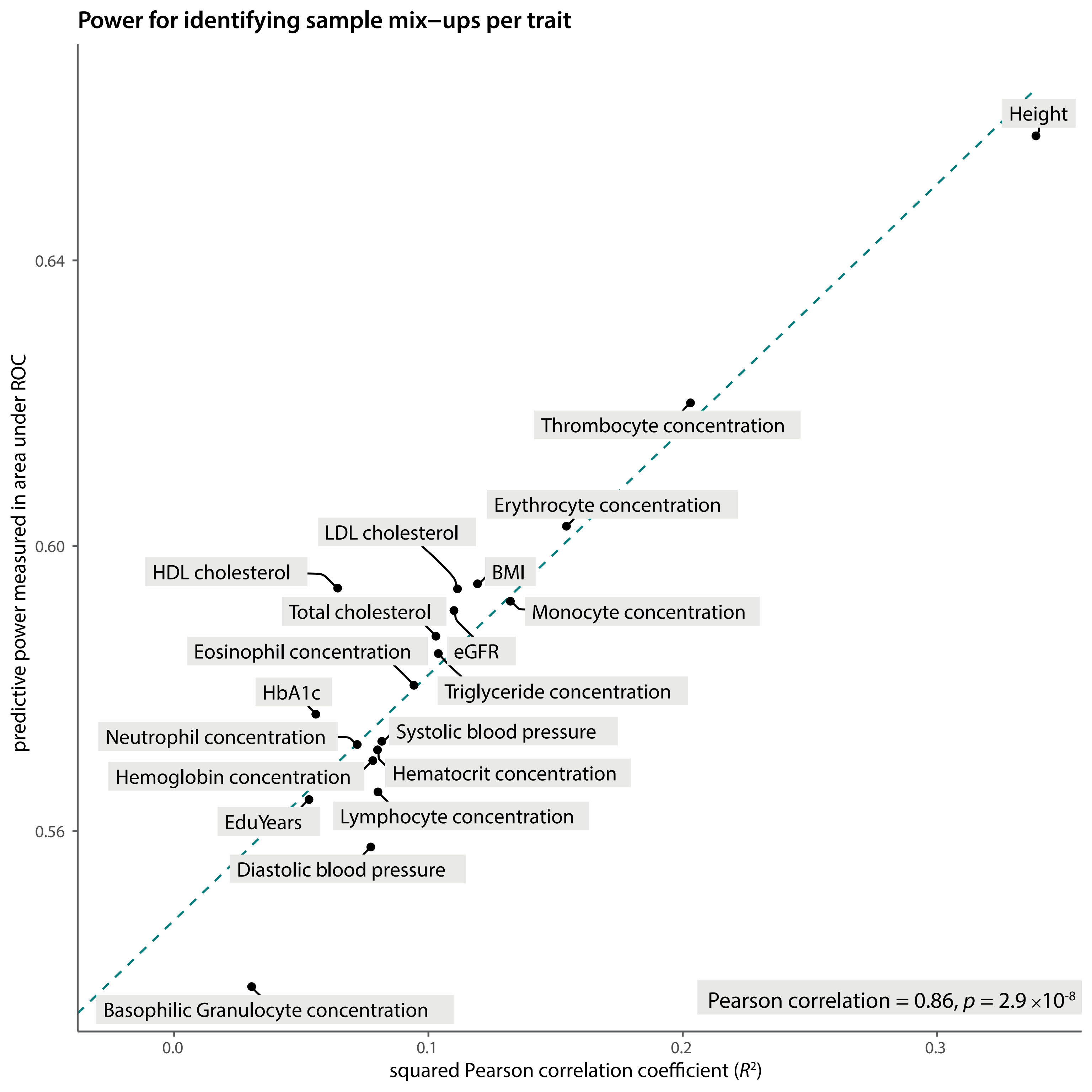

### Supplementary figure 4

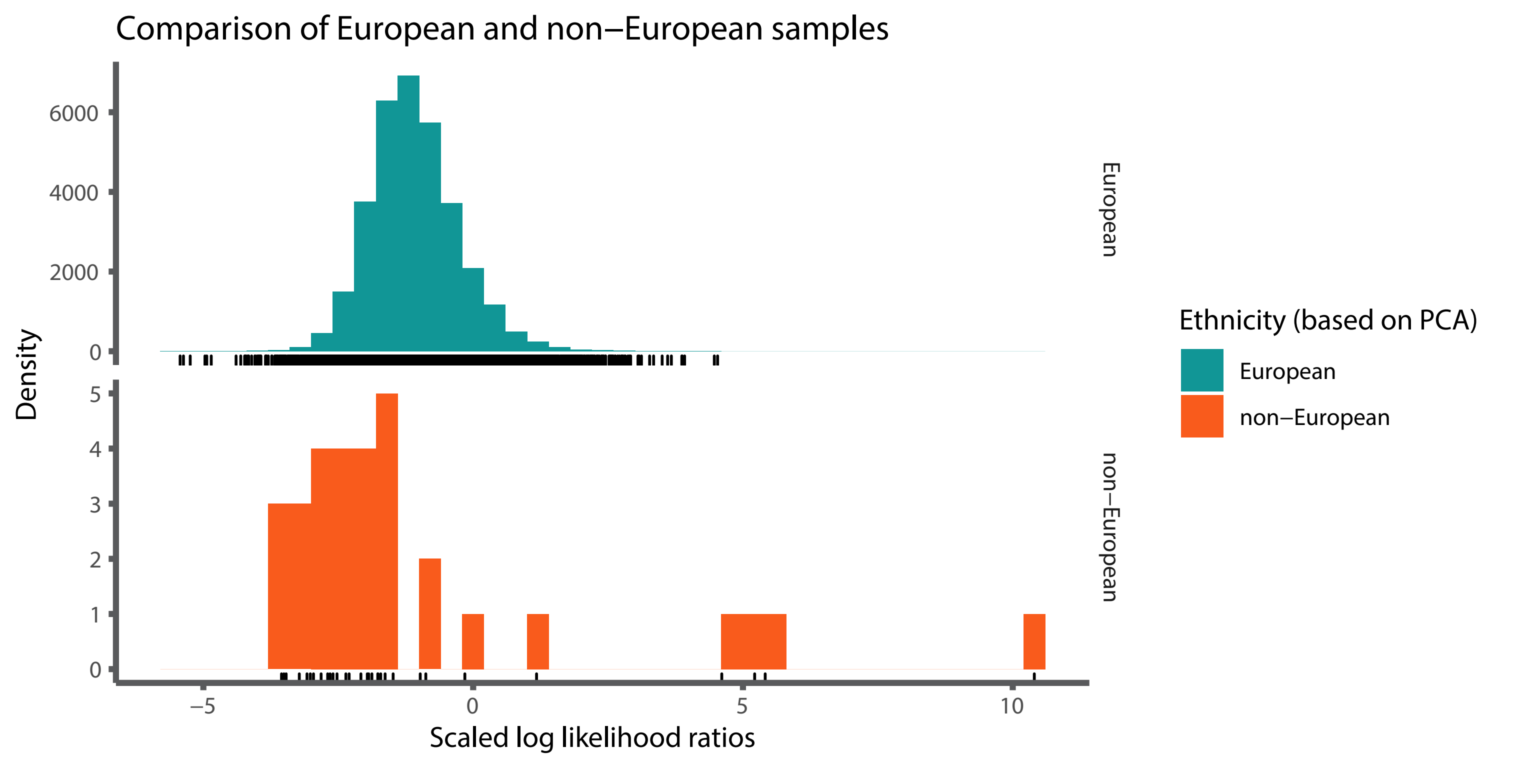

### Supplementary figure 5

#### The basis of mix–up predictions

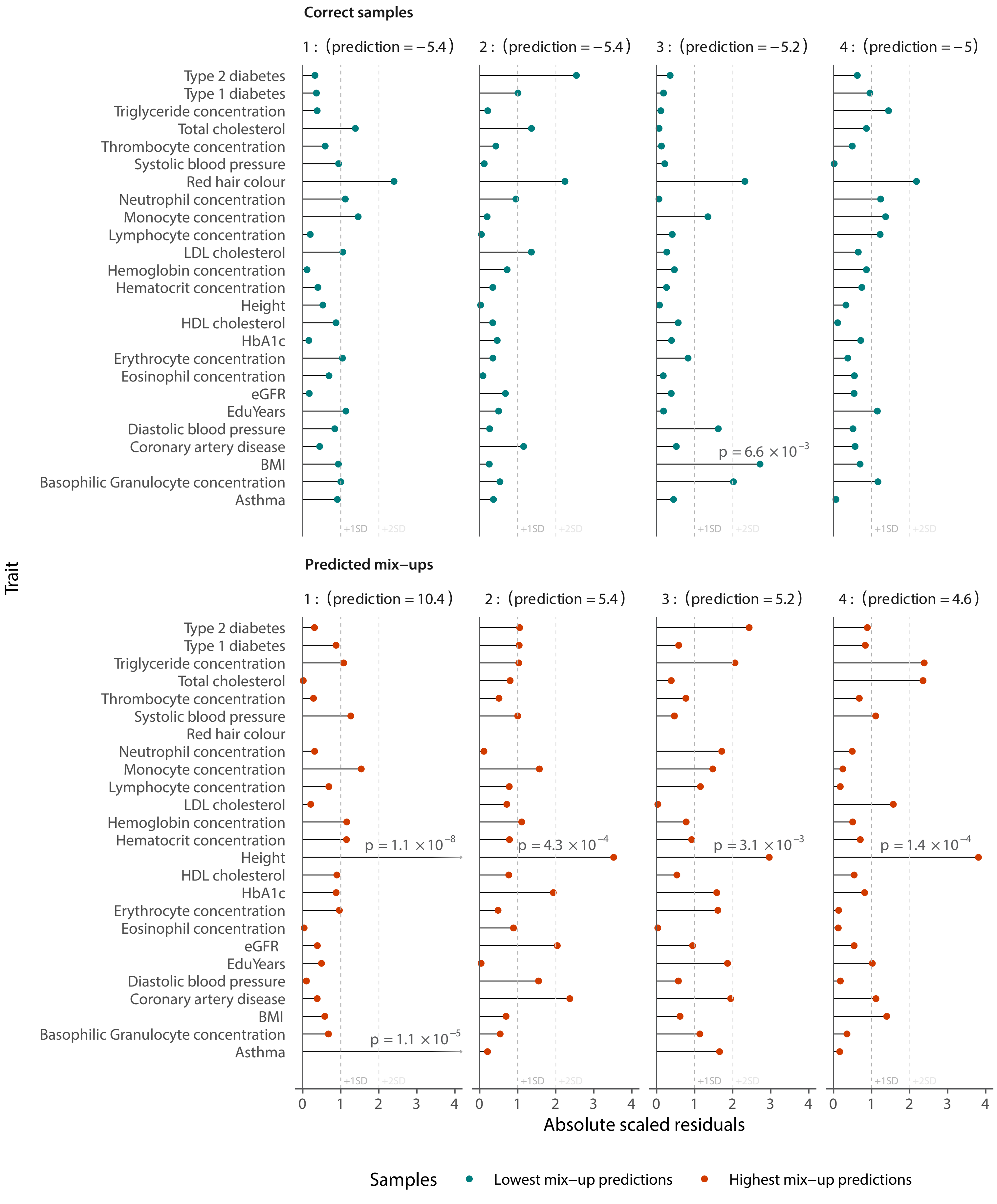
